# Decoding transcriptomic signatures of Cysteine String Protein alpha-mediated synapse maintenance

**DOI:** 10.1101/2023.10.02.560611

**Authors:** Na Wang, Biqing Zhu, Mary Alice Allnutt, Rosalie M. Grijalva, Hongyu Zhao, Sreeganga S. Chandra

## Abstract

Synapse maintenance is essential for generating functional circuitry and decrement in this process is a hallmark of neurodegenerative disease. While we are beginning to understand the basis of synapse formation, much less is known about synapse maintenance *in vivo*. Cysteine string protein α (CSPα), encoded by the *Dnajc5* gene, is a synaptic vesicle chaperone that is necessary for synapse maintenance and linked to neurodegeneration. To investigate the transcriptional changes associated with synapse maintenance, we performed single nucleus transcriptomics on the cortex of young CSPα knockout (KO) mice and littermate controls. Through differential expression and gene ontology analysis, we observed that both neurons and glial cells exhibit unique signatures in CSPα KO brain. Significantly all neurons in CSPα KO brains show strong signatures of repression in synaptic pathways, while upregulating autophagy related genes. Through visualization of synapses and autophagosomes by electron microscopy, we confirmed these alterations especially in inhibitory synapses. By imputing cell-cell interactions, we found that neuron-glia interactions were specifically increased in CSPα KO mice. This was mediated by synaptogenic adhesion molecules, including the classical Neurexin1-Neuroligin 1 pair, suggesting that communication of glial cells with neurons is strengthened in CSPα KO mice in an attempt to achieve synapse maintenance. Together, this study reveals unique cellular and molecular transcriptional changes in CSPα KO cortex and provides new insights into synapse maintenance and neurodegeneration.

**Significance statement:** Synapse maintenance is important for maintaining neuronal circuitry throughout life. However, little is known about molecules that affect synapse maintenance *in vivo*. CSPα, encoded by the *Dnajc5* gene, is a synaptic vesicle chaperone that is linked to synapse maintenance and neurodegeneration. Here, we show by performing single nucleus transcriptomics of CSPα KO cortex that synapse instability is related to repression in synaptic pathways and elevation of autophagy in neurons. However, we find a heterogeneity of glial responses. Additionally, interactions between neurons and glia are increased in CSPα KO, mediated by synaptogenic adhesion molecules. This study provides a novel perspective on into synapse maintenance and reveals unique cellular and molecular transcriptional changes in CSPα KO brains.

## Introduction

Most synapses are stable throughout a lifetime in the healthy adult brain. Of the steps in the lifecycle of a synapse—formation, plasticity, maintenance, and elimination— synapse maintenance is the most poorly understood. However, synapse maintenance is essential for generating functional circuitry and decrement in this process or synapse instability is a hallmark of neurodegenerative diseases (1-3). Therefore, understanding mammalian synapse maintenance pathways at a molecular level is of high biomedical importance.

CSPα is the only chaperone known to be critical for synapse maintenance and prevention of neurodegeneration (4, 5). CSPα is highly conserved across species and is localized to the membrane of synaptic vesicles (SVs) and secretory granules (6). CSPα contains a J domain, typical of Heat shock protein 40 (Hsp40) co-chaperones, through which it activates the ATPase activity of the chaperone Heat Shock Cognate 70 (Hsc70) (7-11) to fold client/substrate proteins (12). CSPα (*Dnajc5*) belongs to the DNAJC class of co-chaperones which are highly selective with a small repertoire of substrates. CSPα chaperones the SNARE SNAP-25 and endocytic GTPase dynamin 1 (13, 14) and regulates the synaptic vesicle (SV) cycle. Hence, CSPα is required for normal neurotransmission and synapse maintenance in the fly (15), worm (16), mouse (5), and human (17). In mice, CSPα deficient animals are relatively normal at birth through postnatal day 21 (P21) but subsequently exhibit altered neurotransmission (5, 18), synapse loss (4), and neurodegeneration. The neurotransmission deficits in CSPα knockout (KO) mice, which are most pronounced in inhibitory neurons, include progressive reduction and desynchronization of synaptic release as well as decreased size of the readily releasable pool of SVs (18, 19). These findings imply that CSPα deficiency leads to progressive synaptic dysfunction and use-dependent loss of nerve terminal integrity. The onset of these phenotypes occurs after weaning in CSPα KO mice, when peak synapse formation is complete, is consistent with a synapse maintenance function.

CSPα has been linked to several neurodegenerative diseases, including genetically to Adult-onset Neuronal Ceroid Lipofuscinoses (ANCL), a lysosomal storage disorder, and mechanistically to Alzheimer’s disease (AD) and Parkinson’s disease (PD) (4, 20-22). Initially, two dominant mutations in CSPα—L115R and L116Δ—were identified as a cause of ANCL (17, 23-25). More recently, C124_C133dup and C128Y mutations have also been described to cause ANCL (26, 27). These ANCL patients exhibit lipofuscin accumulation and profound cortical neurodegeneration leading to presenile dementia. In AD, CSPα expression is downregulated in hippocampus and is suggested to be closely related to early synaptic degeneration and impaired memory formation (20, 28). In the case of PD, CSPα is mechanistically linked to the homeostasis of α-synuclein, a key protein in the pathogenesis of PD (22, 29, 30).

To obtain an accurate and unbiased assessment of the complex cellular and molecular changes associated with synapse instability in the CSPα KO mouse model and to understand cortical degeneration, we performed single nuclei transcriptomics on the cortex of CSPα KO and WT littermate controls at an early stage of synapse loss (postnatal day 24). Interestingly, we observe that both neurons and glial cells exhibit unique signatures that contribute to synapse maintenance. In particular, we detected that a broad range of synapse-related pathways were downregulated, while synaptic autophagy was increased, in all neuronal classes, with more pronounced effects in inhibitory neurons. Furthermore, we inferred that the interactions between neurons and glial cells were increased to maintain synapses, with augmented interaction between the well-known synaptogenic adhesion molecules Neurexin1-Neuroligin1 (Nrxn1-Nlgn1) being the most prominent. Taken together, our data reveal a unique cellular-level view of transcriptional changes in CSPα KO cortex and are a novel resource to investigate mechanisms of synapse maintenance and neurodegeneration.

## Results

### Single nucleus transcriptomic profiling of CSPα KO cortex

Based on our previous analyses of CSPα KO mice at P28 (13) and published work at P50 (31), we hypothesized that P24 represents an early stage of synaptic instability. Therefore, we evaluated the phenotypes of CSPα KO mice at P24 and determined, by electron microscopy (EM), there was modest decrease in synapse number in motor cortex (Fig. 1A, B, WT: 1 ± 0.03; CSPα KO: 0.6 ± 0.03, p < 0.0001). This was due to loss of inhibitory (Type II) synapses in CSPα KO cortex (Fig. 1C, Inhibitory synapse WT: 1 ± 0.04; CSPα KO: 0.5 ± 0.03, p < 0.0001), while excitatory (Type I) synapses did not change (Fig. 1C, Excitatory synapse WT: 1 ± 0.08; CSPα KO: 1.08 ± 0.07, p =0.4761). Additionally, NeuN labelling of neurons in cortex, revealed some neuronal loss in CSPα KO brains (Supplementary Fig. 1A, B). However, these effects were milder than prior analyses at P28 (13) and P50 (31), indicating P24 is an optimal post-weaning age to elucidate the mechanisms that initiate synapse instability in CSPα KO mice.

**Figure 1.**
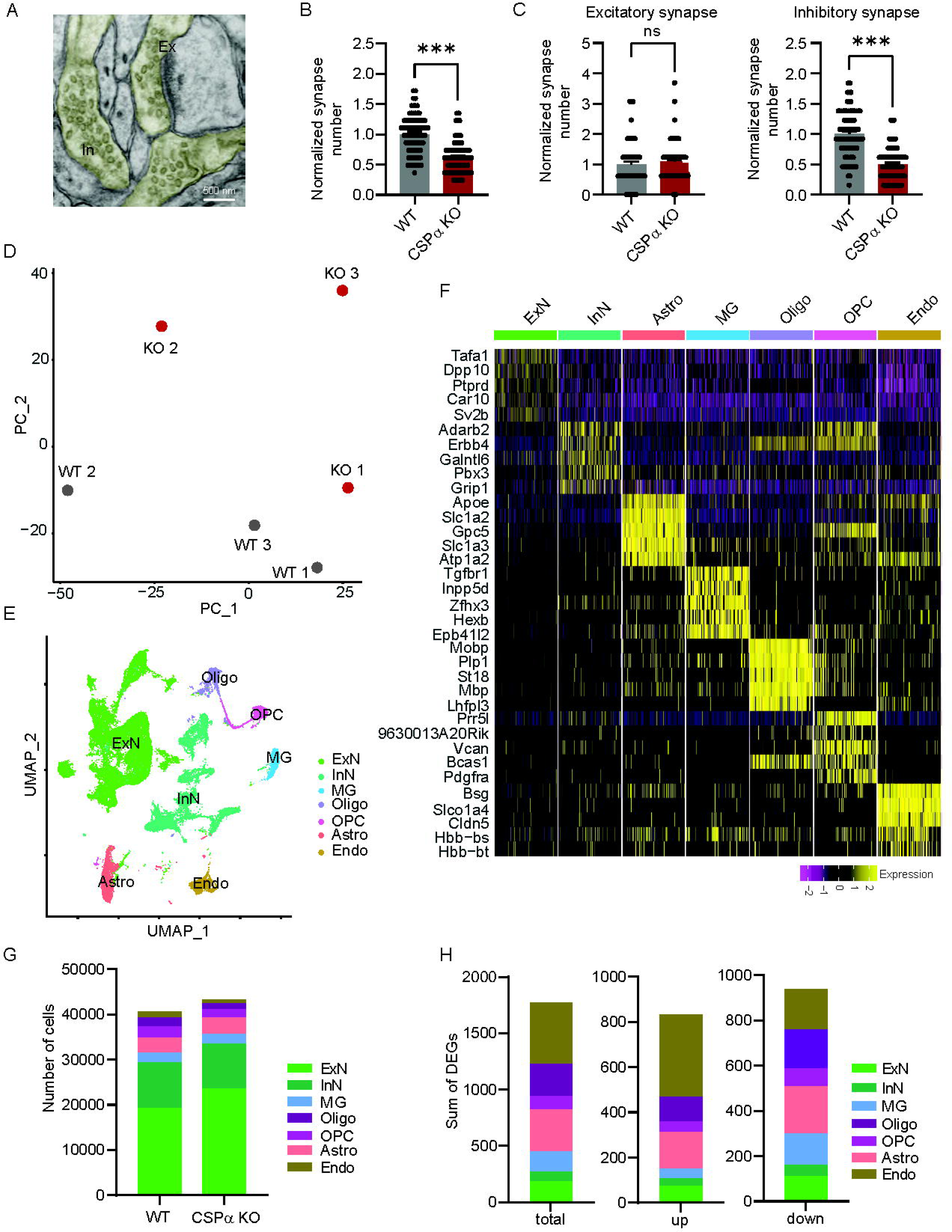
Single-cell transcriptional profiling of the CSPα KO mouse cortex reveals a diversity of cell type changes. **A**. Electron micrograph of excitatory (Ex) and inhibitory (In) synapse in mouse motor cortex. **B.** Quantification of synapse number from electron micrographs of WT and CSPα KO mouse motor cortex. **C**. Quantification of synapse number in WT and CSPα KO brains separated by neurotransmitter type. Note the significantly decreased synapse number of inhibitory synapses but not excitatory synapses in CSPα KO brains. N = 3 brains/genotype with ≥15 micrographs per brain. Each dot represents number of synapses/micrograph. Scale bar: 500 nm. Data are shown as mean values ± SEM after normalization to WT.***p < 0.001, t-test. **D**. PCA for all WT and CSPα KO samples. **E**. UMAP plot of nuclei colored according to the major mouse brain cell types: excitatory (ExN) and inhibitory (InN) neurons, oligodendrocytes (Oligo), astrocytes (Astro), microglia (MG), oligodendrocytes precursor cells (OPC), and endothelial cells (Endo). **F**. Heatmap showing expression of cell type-specific top marker genes in each brain cell type. **G**. Cell numbers of each brain cell type grouped by genotype. **H**. Number of up-regulated, down-regulated, and total differentially expressed genes (DEGs) between WT and CSPα KO for each brain cell type.

To understand CSPα related cellular diversity and transcriptional changes, we performed single nucleus RNA sequencing (snRNA-seq) on the cortex of CSPα KO and littermate wild type (WT) control mice at age P24. We used a mouse brain nuclei isolation protocol to purify nuclei from littermate WT and CSPα KO cortex (n=6; littermates; sex matched) followed by snRNA-seq on the 10x Chromium platform (Supplementary Fig. 2A). After quality control filtering, our snRNA-seq yielded a total of 89,464 high quality nuclei from which we detected a median of 2,000 genes per nucleus (Supplementary Fig. 2B, C, D, E). By Principal Component Analysis, we observe WT and CSPα KO cell distributions largely segregate by genotype (Fig. 1D, Supplementary Fig. 2F). After clustering of nuclei, we identified all the major cortical cell types with known marker genes (32-34): excitatory neurons (ExN), inhibitory neurons (InN), astrocytes (Astro), microglia (MG), oligodendrocytes (Oligo), oligodendrocytes precursor cells (OPC), and endothelial cells (Endo) (Fig. 1E, Supplementary Fig. 3). We observed no large differences in the cell distribution between genotypes (Fig 1G, Supplementary Fig. 2G). We found unique or enriched RNA markers for each cell type by comparing a given cell type to the others by Wilcoxon test: *Tafa1*, *Dpp10*, *Ptprd*, *Car10*, *Sv2b* for ExN; *Adarb2*, *Erbb4*, *Galntl6*, *Pbx3*, *Grip1* for InN; *ApoE*, *Slc1a2*, *Gpc5*, *Slc1a3*, *Atp1a2* for Astro; *Tgfbr1*, *Inpp5d*, *Zfhx3*, *Hexb*, *Epb41I2* for MG; *Mobp*, *Plp1*, *St18*, *Mbp*, *Prr5I* for Oligo; *Lhfpl3*, *9630013A20Rik*, *Vcan*, *Bcas1*, *Pdgfra* for OPC; *Bsg*, *Slco1a4*, *Cldn5, Hbb-bs*, *Hbb-bt* for Endo (Fig. 1F). For both ExN and InN, we identified several subclusters, with 9 ExN and 5 InN subclusters (Supplementary Fig. 2I, J). When we calculated the *Dnajc5* expression value by averaging the expression level in all cells of a given cell type, we found that *Dnajc5* mean expression level was highest in neurons and endothelial cells, with 4-fold higher expression in ExN compared with all glial cells, and 2-fold higher expression in InN (Supplementary Fig. 2H). This finding is consistent with current understanding in the field that CSPα acts cell-autonomously in neurons (Supplementary Fig. 1C, D). Yet, the 1772 differentially expressed genes (DEGs) that were identified between CSPα KO and WT brains were widely distributed in all 7 identified cell types (Fig. 1H), suggesting that both neurons and non-neuronal cells contribute to synapse maintenance.

### CSPα KO neurons exhibit downregulation of synapse organization related pathways

We initially focused on DEGs in neuronal clusters, comparing CSPα KO versus WT. The total number of DEGs is 187 in all ExNs and 82 in all InNs (Fig. 1H). Surprisingly, the majority of DEGs in ExNs and InNs, 112 and 48 respectively, were downregulated, suggesting a signature of neuronal repression (Fig. 2A, B, D, E). We plotted the DEGs as volcano plots for ExNs and InNs to identify prominent changes (Fig. 2A, D). In ExNs, several synaptic related genes, including *Homer1*, *Unc13a*, *Syt7* and *Tmem163* were downregulated (Fig. 2A). Notable increased genes include, *Gria2* (glutamate ionotropic receptor AMPA type subunit 2), the major excitatory neurotransmitter receptor and *Cntn3* (Contactin 3), a neuronal cell adhesion molecule (Fig. 2A). Similarly in InNs, synaptic genes, such as *Nrgn* (Neurogranin) and *Unc13a* were downregulated, while *Grin3a* (glutamate ionotropic receptor NMDA type subunit 3A) was upregulated (Fig. 2D). Next, we examined the pathways enriched in each ExN and InN sub-cluster for down-regulated genes by Gene Ontology (GO) pathway analysis. Intriguingly, the most significantly down-regulated pathway is synapse organization in all ExN sub classes (ExN1-ExN9) (Fig. 2B) and most InNs sub classes (InN2-InN4) (Fig. 2E), supporting our tenet that transcriptional analysis of CSPα KO can yield insights into synapse maintenance pathways. We found that ExN5, ExN6 and InN3 showed the strongest effects (Fig. 2C, F). Prominent DEGs in the synapse organization pathways include ion channels (*Asic2*, *Atp2b2*) and the activity regulated gene *Arc* in ExNs (Table S1), synaptic vesicle associated *Syn1* in InNs (Table S2) and post-synaptic scaffold, *Homer 1* in both ExNs and InNs (Table S1 and Table S2). Interestingly, the genes in synapse organization pathways encode both pre- and post-synaptic proteins suggesting coordinated changes. The other top downregulated pathways in ExNs and InNs were all related to synaptic function (e.g. dendrite development, regulation of synapse structure or activity). Notably, we did not detect any up-regulated pathways in ExNs or InNs, likely because only 75 and 34 upregulated DEGs were detected in ExNs and InNs, respectively (Fig. 1H, Fig. 2A, D). The decreased expression of synaptic genes is congruent with the decreased synapse density exhibited in the CSPα KO cortex *in vivo* and cortical neurons *in vitro* (Fig. 1B, C, Supplementary Fig. 1C, D).

**Figure 2.**
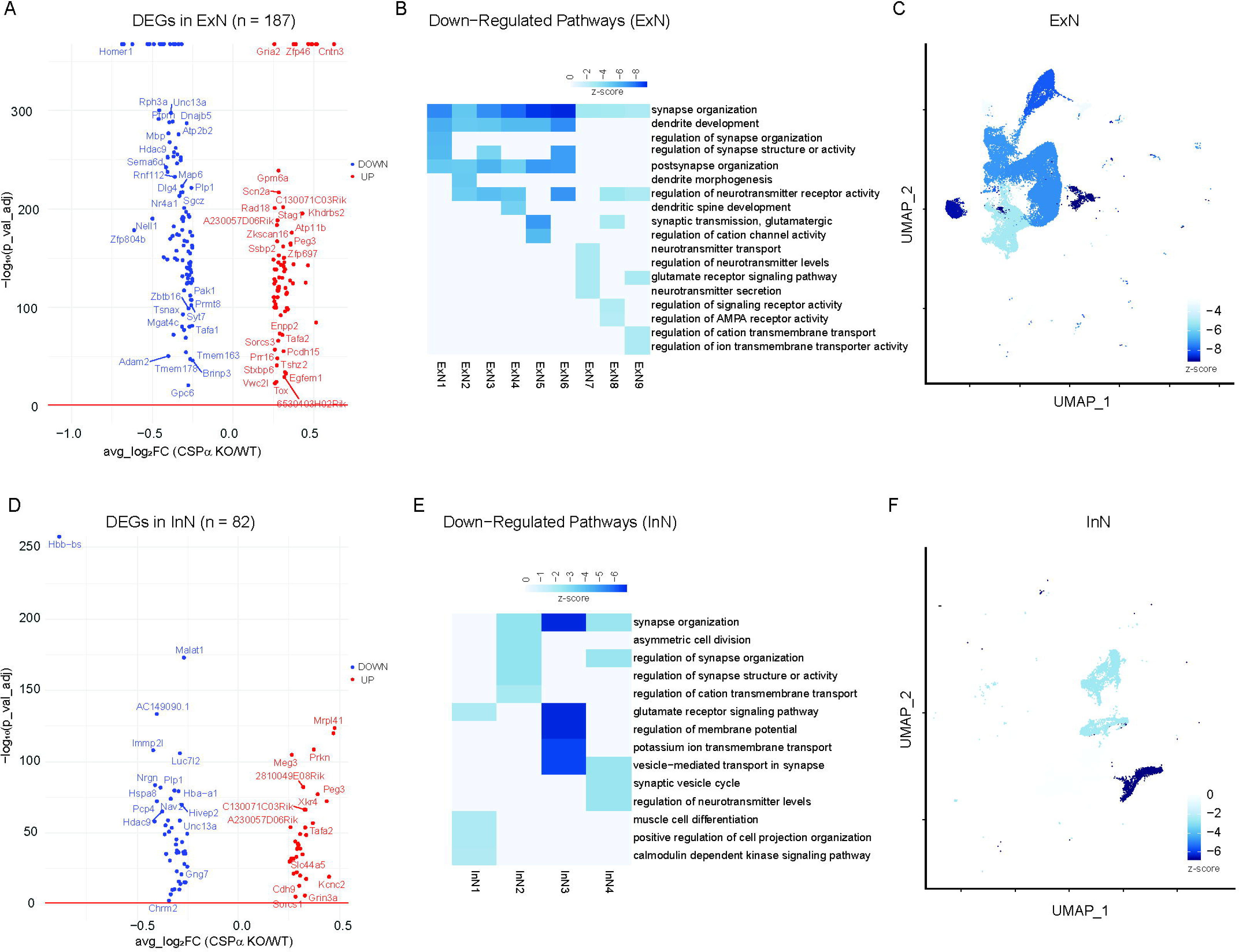
Deletion of CSPα orchestrates neuronal transcriptomic changes in synapse-related genes. **A, D.** Volcano plots for DEGs in ExNs (A) and InNs (D). **B, E**. Gene ontology (GO) pathway analysis of DEGs between WT and CSPα for downregulated pathways in ExNs (B) and InNs (E). Scale: z-score of pathway p-values. **C, F**. UMAP plot of the Synapse Organization pathway in ExN and InN subclusters. Scale: z-score of pathway p-values.

To evaluate the role of CSPα’s chaperone activity in the observed synaptic alterations, we tested the expression level of CSPα interactors such as other heat shock proteins/chaperones and known substrates including Snap25, Dnm1, and Sept5 (Supplementary Fig. 4A, D). *Hspa8*, which encodes for Hsc70, the main partner of CSPα, was downregulated in both CSPα KO ExNs and InNs (Supplementary Fig. 4A). We determined that decreased *Hspa8* gene expression leads to lower Hsc70 protein levels by western blotting of WT and CSPα KO synaptosomes (Supplementary Fig. 4B, C, WT: 1 ± 0.05; CSPα KO: 0.74 ± 0.08, p = 0.047). Additionally, we found distinct sets of heat shock genes were downregulated in CSPα KO neurons. In ExNs, *Hsph1*, *Dnaja4*, *Hsp90ab1* were downregulated (Supplementary Fig. 4A), while in InNs, Hsph1, *Hspd1*, *Dnaja4*, *Hsp90aa1*, *Chordc1*, *Dnaja1*, and *Hsp90ab1* were downregulated (Supplementary Fig. 4A). Collectively, this indicates a decrement of chaperone capacity in neurons upon deletion of CSPα. The transcript levels of known CSPα substrates *Snap25*, *Dnm1*, and *Sept5*, were largely unaltered in both ExNs and InNs (Supplementary Fig. 4D). Congruently, we did not detect any significant difference in the expression of *Dnm1* and *Sept5* by qPCR in CSPα KO cortex (Supplementary Fig. 4E; *Sept5*: WT: 0.99 ± 0.02; CSPα KO: 0.90 ± 0.08; p = 0.3125; *Dmn1*: WT = 1.00 ± 0.05; CSPα KO= 1.17 ± 0.09; p = 0.26). Together, this suggests the effects of CSPα deletion impacts the transcription of other chaperones but not its substrates, yet chaperone levels will impact substrate protein folding.

Next, we analyzed upregulated genes in ExNs and InNs individually as we did not find any upregulated pathways. The expression of critical autophagy associated genes was increased in both ExNs and InNs (Fig. 3A, B), including the SV associated protein Atg9a. Recently, Atg9a was demonstrated to couple the SV cycle to presynaptic autophagy (35) and is well-poised to coordinate with CSPα. However, it is not clear whether Atg9a has any relationship with CSPα or contributes to the phenotypes of CSPα KO mice. We performed immunocytochemistry on WT and CSPα KO cultured cortical neurons and confirmed that Atg9a was partially colocalized with CSPα in WT neurons (Fig. 3C, D; 56.7% colocalization), indicating a presynaptic localization. Comparing the distribution of Atg9a in WT and CSPα KO neurons, we find the colocalization of Atg9a and Amphiphysin 2, a presynaptic endocytic protein, was significantly increased in CSPα KO neurons (Fig. 3E, F; WT: 24.98% ± 1.39%, CSPα KO: 53.04% ± 8.09%, p = 0.009), consistent with the possibility of increased autophagosome biogenesis in CSPα KO brains. To determine whether autophagy was indeed increased in CSPα KO neurons, in line with the transcriptomic changes, we performed EM on WT and CSPα KO motor cortex (n=3 mice/genotype; P24). We quantified the number of presynaptic autophagosomes and categorized them by synapse type. We observe a significant increase in autophagosomes in CSPα KO inhibitory synapses (Fig. 3G, I, WT: 0.03 ± 0.01; CSPα KO: 0.21 ± 0.03, p< 0.0001) with a trend in excitatory synapses (Fig. 3H, WT: 0.03 ± 0.02; CSPα KO: 0.07 ± 0.02, p=0.28). Since neurons exhibit compartment – specific autophagy, which depends on polarized structures and rapid autophagy flux (36), it is possible for excitatory synapses not to show increased autophagosomes, even though the expression level of autophagy genes was increased. Intriguingly, many of the autophagosomes appear to contain SVs or SV-like vesicles. To investigate whether presynaptic autophagy is involved in SV degradation, we quantified autophagosomes containing SVs or SV-like vesicles in WT and CSPα KO cortex. We also determined the diameter of cargo vesicles. We found that the percentage of autophagosomes containing SV-like structures was increased significantly in CSPα KO inhibitory synapses (Fig. 3G, J, WT: 2.3% ± 1.6%; CSPα KO: 14.1% ± 3.6%, p=0.0025). Furthermore, the diameters of the SV-like structures inside the autophagosomes were comparable to SVs outside the autophagosomes (Fig. 3G, K, Inside: 46.58 ± 4.30; Outside: 38.96 ± 1.94, p= 0.08). We previously demonstrated that SV numbers in CSPα KO neurons were reduced (13), and our current results indicate that their removal may occur partially through autophagy.

**Figure 3.**
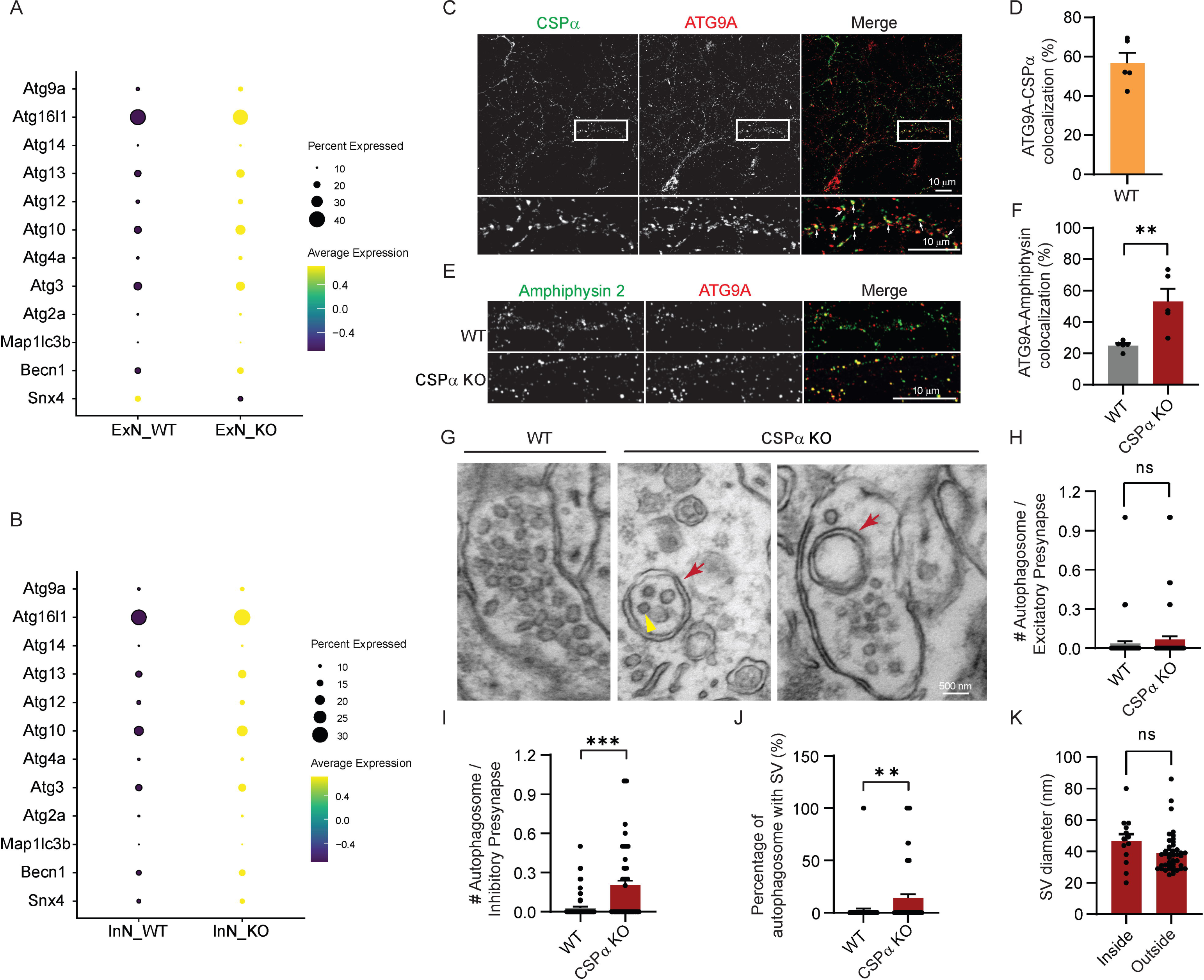
Presynaptic autophagy is increased in CSPα KO brains. **A**. Dot plot for expression of autophagy related genes in WT and CSPα KO ExNs. **B**. Dot plot for expression of autophagy related genes in WT and CSPα KO InNs. **C.** ATG9a partially colocalizes with CSPα in cortical neurons. Scale Bar: 10 μm. **D**. Percent of ATG9a puncta co-localized with CSPα. **E**. The distribution of ATG9a and amphiphysin 2 in WT and CSPα KO cortical neurons. Scale Bar: 10 μm. **F**. Quantification of the ATG9a-Amphiphysin 2 colocalization. **G**. Representative EM micrographs from WT and CSPα KO M1 motor cortex, Age=P24, n=3 brains/genotype, ≥15 micrographs/brain. Red arrows indicate the double-membraned autophagosomes. Yellow arrows indicate SV-like structures. **H**. Quantification of the number of autophagosomes in each excitatory presynapse. **I.** Quantification of the number of autophagosomes in each inhibitory presynapse. **J**. The percentage of autophagosomes containing SV-like structures. CSPα KO synapses contain significantly more autophagosomes with SV-like structures compared with WT synapses. **K**. Quantification of the diameters of SV-like structures inside autophagosomes or SVs outside autophagosomes. Note diameter of SV-like structures inside autophagosomes and diameter of SVs are similar. Data are presented as mean ± SEM. Scale Bar: 500 nm. **p < 0.01, ***p < 0.001, t-test.

### Glial cells exhibit heterogenous signatures of synapse maintenance

We examined the DEGs in all non-neuronal cell types—MG, Astro, Oligos, OPCs, and Endo cells. Similar to neurons, DEG analysis revealed that the majority of DEGs in all glial cell types were down-regulated (MG: 141 down-regulated DEGs and 42 up-regulated DEGs; Astro: 208 down-regulated DEGs and 162 up-regulated DEGs; Oligos: 174 down-regulated DEGs and 110 up-regulated DEGs; OPCs: 77 down-regulated DEGs and 45 up-regulated DEGs; Endo cells: 178 down-regulated DEGs and 366 up-regulated DEGs) as seen by the volcano plots (Fig. 1H, Supplementary Fig. 5). The most prominent down-regulated DEG in MG was Malat 1 which is a critical regulator of CD8^+^ T cell differentiation (Supplementary Fig. 5A). ApoE, a well-known gene that influences Alzheimer’s risk, was upregulated in both MG and Astro (Supplementary Fig. 5A, B). Similar to our analysis for neurons, we then performed pathway enrichment for both down-regulated and up-regulated DEGs in each cell type by GO pathway analysis. Interestingly, unlike neurons, we could identify upregulated and downregulated pathways in all glia cell types (Fig. 4 A, B). MG were enriched for the regulation of immune effector process (Fig. 4A) which suggested MG may be involved in immune responses in CSPα KO mice. The upregulated pathways in Astro contained small GTPase signal transduction and Ras protein signal transduction (Fig. 4A) which are important for cell-cell adhesion and other processes that involve the regulation of cytoskeletal dynamics (37). In the down regulated pathway analysis, we detected synaptic-related pathways such as synapse organization, axonogenesis, and myelination in MG and Oligos (Fig. 4B). Taken together, the upregulated and downregulated pathways in glia cells suggested that they exhibit distinct signatures upon removal of CSPα.

**Figure 4.**
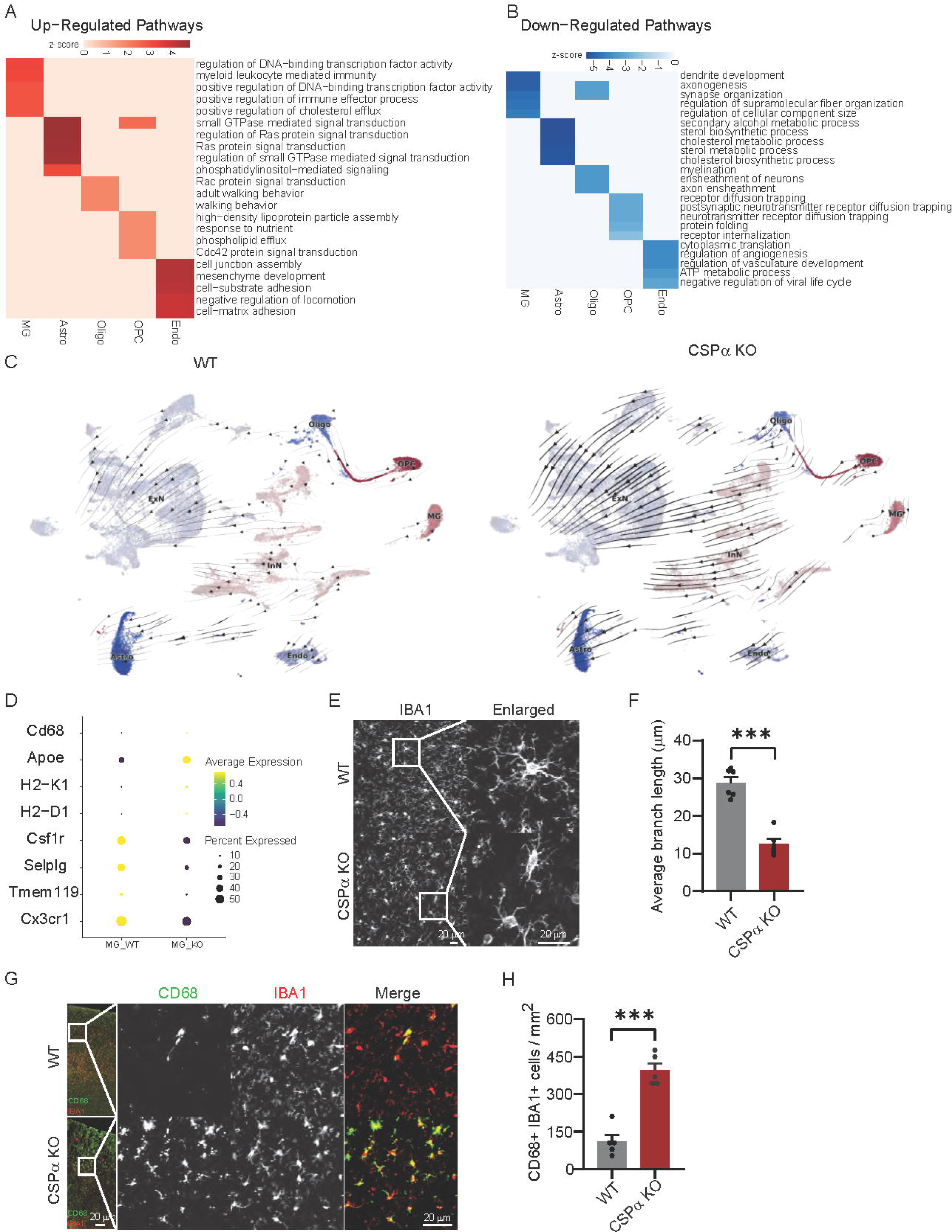
Glial cell signatures indicate they play diverse roles in synapse maintenance. **A, B.** GO pathway analysis of DEGs between WT and CSPα for (A) up-regulated and (B) down-regulated pathways in each non-neuronal cell type. Scale: z-scores of pathway p-values. **C**. RNA velocity analysis for each cell type as a function of genotype. **D**. Targeted analysis of key MG genes. **E**. The morphology of MG in WT and CSPα KO cortex revealed by IBA1 staining. **F**. Quantification of the branch length for MG, n=6 brains/genotype. **G.** CD68 and IBA1 staining of MG in WT and CSPα cortex. **H.** Quantification of the number of CD68 and IBA1 double positive MG, n=5 brains/genotype. Data are presented as mean ± SEM. Scale Bars: 20 μm. ***p < 0.001, t-test.

To analyze expression dynamics, we mapped RNA velocity (38) in CSPα KO and WT cortex. Overlaying these trajectories on the UMAP (Fig. 1E, Supplementary Fig. 2I), we find trajectory shifts in MG, OPC, ExN4, ExN7, InN2, InN3 and InN4 between the two genotypes (Fig. 4C). Since the trajectories of MG were altered dramatically compared to other cell types, we further investigated additional targeted DEGs in MG focusing on homeostatic and disease associated microglia (DAM) genes. Interestingly, homeostatic genes, such as *Cx3cr1*, *Tmem119*, *Selplg*, and *Csf1r* were downregulated in CSPα KO MG, while DAM genes including *H2-D1*, *H2-K1*, and *ApoE* were notably upregulated in CSPα KO cortex compared to WT (Fig. 4D, Supplementary Fig. 5A).

As MG exhibited unique signatures in GO and RNA velocity analysis, we stained WT and CSPα KO cortex for MG with an IBA1 antibody to characterize the morphology of microglia. Imaging revealed that microglia morphology was dramatically altered in CSPα KO cortex in comparison to WT mice (Fig. 4E). We quantified microglia branch length and found they were significantly reduced in CSPα KO cortex (Fig. 4F, WT: 28.77 ± 1.53 µm; CSPα KO: 12.59 ± 1.27 µm, p < 0.0001). To assess microglial activation, IBA1 and CD68 were co-stained to label all microglia and activated microglia, respectively (39). Notably, we found microglia had significantly increased CD68 immunoreactivity in CSPα KO cortex (Fig. 4G, H, WT: 110.9 ± 26.9; CSPα KO: 396.0 ± 27.7, p < 0.0001). This result was consistent with the increased RNA expression of CD68 in CSPα KO microglia (Fig. 4D).

### The increased interaction probability of Neurexin1-Neuroligin 1 in CSPα KO neurons and **glial cells**

Since we detected some synaptic-related pathways were also downregulated in certain glial cells (Fig. 4B; Fig. S5), we wanted to understand neuron-glia relationships in WT and CSPα KO brains. We imputed the inter-cellular communication network using CellChatDB (40), an analysis tool that can quantitatively characterize and compare inferred inter-cellular communications to enable identification of specific signaling roles played by each cell population. We first tested interactions between Astro and InNs and found increased interaction strength between these two cell types in CSPα KO brains (Supplementary Fig. 6A). We next identified ligand-receptor interactions from Astro to InNs driving this increased interaction and found the strongest and consistently increased pair was Neurexin 1-Neuroligin 1 (Nrxn1-Nlgn1) (Fig. 5A). Nrxn1 and Nlgn1 have been extensively studied as synaptic cell adhesion molecules that connect pre- and postsynaptic neurons and mediate signaling across the synapse (41, 42), which modulates synaptic activity and determines the properties of neuronal networks (43). However, recent studies suggest a role for Nrxn1 and Nlgn1 in glia in mediating synapse formation and function (44, 45). Notably, the other ligand-receptor pairs, driving the increased interactions between Astro and InNs, were also synaptic adhesion molecules including other neurexins and immunoglobulin super family adhesion molecules such as Cadm1(46) (Fig. 5A). To our surprise, this pattern of higher synapse adhesion molecular interactions was seen between other glial cells (MG, Oligo, OPC) and InNs as well as ExNs (Fig. 5B, Supplementary Fig. 6B-G). In each case, the Nrxn1-Nlgn1 pair was the most prominent ligand-receptor pair (Fig. 5, Supplementary Fig. 6). It is noteworthy that we did not observe ligand-receptor pairs important for synapse elimination such as complement.

**Figure 5.**
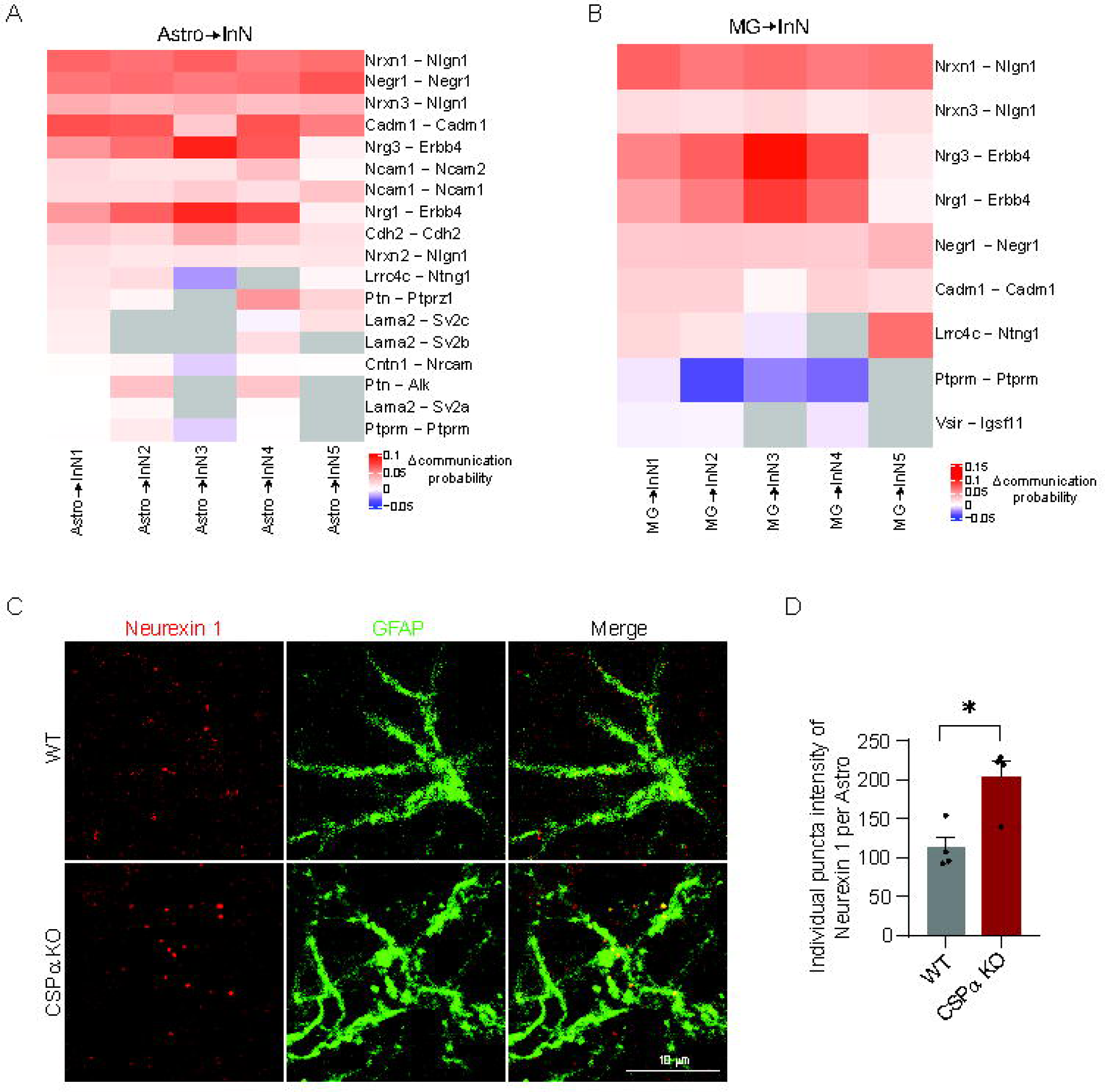
Receptor-ligand interaction pair alterations in CSPα KO mice. **A.** The receptor-ligand pairs exhibiting increased interaction between Astro and InNs in CSPα KO brains as imputed by CellChatDB. Scale: Communication probability of CSPα KO brains – communication probability of WT brains. **B**. The receptor-ligand pairs exhibiting increased interaction between MG and InNs in CSPα KO mice. Note that Nrxn1-Nlgn1 was a top pair for glia-InN interactions. Scale: Communication probability of CSPα KO brains – communication probability of WT brains. **C.** Neurexin 1 and GFAP staining in WT and CSPα cortex. **D.** Quantification of intensity of individual puncta of Nrxn1 on each GFAP positive astrocyte, n=4 brains/genotype. Data are presented as mean ± SEM. Scale Bars: 10 μm. *p < 0.05, t-test.

To test these findings, we stained for Astro using the marker GFAP and Nrxn 1 (Fig. 5C) and quantified the individual puncta intensity of Nrxn1 on each Astro. We found that the individual puncta intensity of Nrxn1 was significantly increased in CSPα KO Astro (Fig. 5D, WT: 112.3 ± 14.2; CSPα KO: 202.8 ± 21.14, p = 0.012). We were unable to stain for Nlgn1 due to poor antibodies. Taken together, our results suggest that in CSPα KO brains, synapse adhesion molecular interactions, typically used for synapse formation between neurons, are utilized for glial-neuronal communication to regulate synapse maintenance.

## Discussion

CSPα is an essential molecular co-chaperone that facilitates synapse maintenance. Here, we systematically characterized the transcriptomic changes in the cortex of WT and CSPα KO mice and utilized the gained insights to experimentally validate mechanisms underlying CSPα-dependent synapse maintenance. We identified that deletion of CSPα lead to downregulation of a broad range of synapse-related pathways in neurons, which was accompanied by upregulation of synaptic autophagy, with these effects being more pronounced in inhibitory neurons. Our data are also suggestive of a diversity in glial responses--both reactive and supportive--in synapse maintenance. Intriguingly, we found enhanced interaction between glia and neurons mediated by synaptic adhesion molecules, suggesting that synapse maintenance is not just a lack of synapse elimination. The systematic and unbiased nature of our approach enabled us to illuminate novel aspects of synapse maintenance. We encourage the scientific research community to utilize this rich dataset as a resource for generating hypothesis about neuronal chaperones, glial roles, structural plasticity, and synapse loss in neurodegeneration (NCBI GEO XXX).

### CSPα deficiency and synaptic dysfunction

To understand the contribution of CSPα’s co-chaperone function to the downregulation of synapse related pathways in CSPα KO neurons, we examined CSPα’s interactors and substrates (47-49). CSPα binds Hsc70 (Gene: *Hspa8*) and together they refold client proteins such as SNAP25 and dynamin 1 (14, 31). SNAP25 is required for the assembly of the SNARE (soluble N-ethylmaleimide-sensitive factor attachment receptor) complex, which is essential for exocytosis and neurotransmitter release, while dynamin 1 is a GTPase involved in clathrin mediated endocytosis, thus CSPα is important for exo-endocytosis coupling. Interestingly, Hspa8 was one of the genes involved in the most downregulated pathway of ‘Synapse Organization’ (Table S1, S2). We determined that protein expression level of Hsc70 was also reduced in synaptosomes (Supplementary Fig. 4B, C). Notably, by examining other chaperones in ExNs and InNs, we observed strong signature of repression of many classes of chaperones (Supplementary Fig. 4A) indicating a wide-spread chaperone deficiency and proteostasis collapse in neurons. How chaperones regulate synaptic protein transcription remains to be addressed and is an interesting line of investigation to further understand synapse maintenance. The transcript levels of CSPα clients were unchanged (Supplementary Fig. 4E), as we have shown previously (13), supporting that substrate interactions only occur at the protein level. Collectively, our data indicate that CSPα affects synaptic transcription through expression and function of its chaperone partners.

### Dysregulation of synaptic autophagy

In neurons, there is mounting evidence suggesting that autophagy is initiated at presynaptic termini. For instance, autophagic vesicles (AVs) appear in presynaptic boutons in response to rapamycin and enhanced Sonic hedgehog signaling (50, 51), and they appear in axons after synaptic activity (52). Studies in model organisms support the importance of autophagy and autophagic machinery in early synaptic development (53, 54). We identified upregulated autophagy-related genes in CSPα KO neurons (Fig. 3A, B) and increased autophagosomes in CSPα KO inhibitory synapses by ultrastructural evaluation (Fig. 3G, H, I). More interestingly, some of the autophagosomes contain SV-like structures (Fig. 3G, J), and the diameters of these SV-like structures are similar to the SVs outside the autophagosomes (Fig. 3G, K). These results suggest that lack of CSPα dysregulates the SV cycle, most likely at the exo-endo coupling step, leads to improper SVs that are removed by synaptic autophagy. A prominent candidate gene that couples the SV cycling and autophagy is *Atg9a* (35). Indeed, we find that the co-localization of Atg9a and Amphiphysin 2 in presynapses in CSPα KO is significantly increased (Fig. 3E, F). Live imaging experiments will be needed to clarify if Atg9 is preferentially sorted to SVs in CSPα KO synapses and if this initiates removal of SVs. We aim to tackle these questions in future studies. Our results highlight that presynaptic proteostasis and autophagy are connected processes and that CSPα plays a prominent role.

### Role of Glia in CSPα KO phenotypes

We observed diverse glial responses to the loss of CSPα. As CSPα expression is 2-4 fold higher in neurons than in any glial cell type (Supplementary Fig. 2H), these are likely driven by responses to the extensive synaptic dysfunction observed in CSPα KO neurons. In MG, up-regulation of pathways such as myeloid leukocyte mediated immunity and positive regulation of immune effector process, as well as RNA trajectory analysis (Fig. 4A, C) suggest that MG are mediating an immune process in CSPα KO mice. MG show broad signatures of activation, including decreased branch length (Fig. 4E, F) and increased CD68 puncta (Fig. 4G, H). While morphology and CD68 positivity signify broad nonspecific activation of MG, the upregulation of DAM genes and downregulation of homeostatic gene expression (Fig. 4D) in MG indicate that these cells are responding to neuronal damage signals (55). The downregulation of pathways related to synapse organization and axonogenesis further reaffirm this changing role of MG in CSPα KO mice (Fig. 4B). While MG appear to have adopted a DAM phenotype in CSPα KO cortex by P24, the response of Astro to this dysfunction is less clear. Gfap is upregulated in Astro (Supplementary Fig. 5B) consistent with an increase in reactive astrocytes (56). However, we also observe several upregulated pathways in Astro related to cell-cell adhesion, such as small GTPase signal transduction and Ras protein signal transduction (Fig. 4A), which may indicate a supportive role for astrocytes in maintaining synapses (37). Although Nrxn1 expression is downregulated in CSPα KO Astro compared to WT (Supplementary Fig. 5B), the communication probability of Astro-InN Nrxn1-Nlgn1 is significantly increased in CSPKO vs WT (Fig. 5B). Our immunostaining showed an increased puncta intensity of Nrxn1 per astrocyte in KO cortex (Fig. 5D), which may be consistent with this overall increased probability of Nrxn1-Nlgn1 interaction. So, in both MG and Astro, we identify reactive signatures but increased probability of synapse adhesion interactions with neurons, which might counteract their reactive roles at this early stage of synapse instability in CSPα KO mice.

### Role of synaptic cell- adhesion molecules in synapse maintenance

Synaptic cell-adhesion molecules are transmembrane or membrane-tethered proteins involved in the formation and function of synapses. They belong to several protein families, including Nrxns, Nlgns, cadherins, protocadherins (Pcdhs), immunoglobulin superfamily (IgSF) proteins, and leucine-rich repeat (LRR) proteins (57). Nrxns and Nlgns are the best characterized synaptic cell adhesion molecules (58, 59) and they play a role during the very initial steps of synaptogenesis through neuron-neuron interactions (60-62). However, analysis of mice lacking these cell adhesion molecules surprisingly revealed that Nlgns and Nrxns are essential for synaptic function, not necessarily synapse formation (63-65). These data suggest roles of Nrxns and Nlgns and other synapse cell-adhesion molecules in both structural and functional synapse maintenance (66). Indeed, adult deletion of these genes supports this tenet (67, 68). In our results, we surprisingly identified Nrxn1-Nlgn1 as the receptor-ligand pair that is increased and mediates glial-neuron interactions (Fig. 5; Supplementary Fig. 6) in CSPα KO mice. Our data adds to an emerging role of Nrxns and Nlgns in glia, not just neurons, contributing to synapse formation and maintenance (44, 45). We presently do not know the purpose to the higher glial-neuron Nrxn1-Nlgn1 interactions in CSPα KO mice. We speculate that this a compensatory mechanism which has revealed that synapse maintenance is an active, ongoing process mediated, in part, by synapse adhesion molecules throughout the lifespan of a synapse. How synaptic cycling, chaperone activity, and synapse adhesion molecules are coordinated to achieve synapse stability is an important question that remains to be addressed.

## Materials and Methods

### Animals

All animal procedures were performed in accordance with the National Institutes of Health Guide for Care and Use of Experimental Animals and were approved by the Yale University Institutional Animal Care and Use Committee. CSPα KO mice have been previously described (5) and are available from Jackson Laboratories (#006392). CSPα KO mice in our colony were made congenic by >10 breedings to C57BL/6J mice. All mice used for experiments came from heterozygous breedings and we used littermate CSPα KO and WT mice on a pure C57BL/6J background. Male and female mice were used in equal numbers for biochemical and sequencing analyses. Mice were maintained in a 12-hour light/dark cycle with standard mouse chow and water *ad libitum*.

### Nuclei isolation from WT and CSPα KO cortex

Freshly dissected mouse cortex from P24 littermate mice were gently Dounce homogenized in 2 ml of ice-cold Nuclei EZ Prep buffer (Sigma, Cat.NUC101-1KT) with a loose pestle, and then with a tight pestle each for 25 times. The resulting homogenate was incubated on ice for five minutes with an additional 2 mL of cold EZ Prep buffer. Nuclei were centrifuged at 500xg for five minutes at 4°C. Pelleted nuclei were then washed in 4 mL of cold EZ Prep buffer, incubated on ice at five minutes, and centrifuged again at 500xg for five minutes at 4°C. Pelleted nuclei were then washed in 4 mL of Nuclei Suspension Buffer containing 0.2% RNase inhibitor (Clontech/Takara, Cat. 2313B) and centrifuged at 500xg for five minutes at 4°C. Finally, nuclei were resuspended in 500μl of Nuclei Suspension Buffer, and counted using a Countess II FL Cell Counter (Applied Biosystems, Cat. A27978). Single-nuclei suspensions were diluted to approximately 1000 nuclei per μl in Nuclei Suspension Buffer.

### Droplet-based single nucleus RNA sequencing and data alignment

snRNA-seq libraries were prepared by the Chromium Single Cell 3 ′ Reagent Kit v3 chemistry according to the manufacturer’s instructions (10x Genomics). The generated snRNA-seq libraries were sequenced using Illumina NovaSeq6000 S4 at a sequencing depth of 300 million reads per sample. For snRNA-seq of mouse cortex, a custom pre-mRNA mouse genome reference was generated with Mouse mm10 (2020-A) that included pre-mRNA sequences, and snRNA-seq data were aligned to this Mouse mm10 (2020-A) reference to map both unspliced pre-mRNA and mature mRNA using CellRanger version 6.0.1. The raw data are available on NCBI GEO (Accession XXXX).

### Single cell quality control and clustering

For quality control, nuclei with mitochondrial content >5% were removed. Additionally, nuclei with less than 200 genes (poor quality nuclei) and more than 7,000 genes (potential doublets) per nucleus were eliminated. Seurat (version 4.0.2) single cell analysis R package was used for processing the snRNA-seq data, followed by the integration of all samples and dimensionality reduction using principal components analysis (PCA). Then, Uniform Manifold Approximation and Projection for Dimension Reduction (UMAP) was applied to visualize all cell clusters, and the classification and annotation of distinct cell types was based on known marker genes of each major brain cell type.

### Differential expression genes (DEGs) analysis

Differential expression genes analysis for snRNA-seq data was performed using the Wilcoxon Rank Sum test with the function FindMarkers of the Seurat package (4.0.5) in R. The genes were filtered using the default parameters (32, 69).

### Gene ontology (GO) pathway analysis

Gene-set enrichment analysis was performed using the function enrichGO from the R package clusterProfiler. GO terms from biological process (BP), cellular component (CC), and molecular function (MF) subontologies were considered (32, 69).

### Electron microscopy

CSPα KO mice and their WT littermate controls at P24 (n=3/group; sex-balanced) were anaesthetized using isoflurane inhalation and perfused intracardially using 2% PFA and 2% glutaraldehyde prepared in 0.1M PB. Brains were harvested and immersed, followed by overnight in 0.1 M cacodylate buffer with 2.5% of glutaraldehyde and 2% PFA. M1 motor cortex was dissected, and further processed at the Yale Center for Cellular and Molecular Imaging, Electron Microscopy Facility following previously published protocols (70). EM imaging was performed using FEI Tecnai G2 Spirit BioTwin Electron Microscope. Images were analyzed blinded to the genotype using FIJI software for synapse number and ultrastructural analysis. Type I or excitatory synapses and Type II or inhibitory synapses were identified by morphological features of the postsynaptic density.

### Immunohistochemistry

Immunohistochemistry on P24 CSPα KO and WT control brains (n=5-6/group; sex-balanced) was performed using paraformaldehyde fixed coronal sections, as we have done previously (71). Primary antibodies used for this study are: CSPα (Chemicon, Cat# AB1576; RRID: AB_90794), NeuN (Millipore, Cat# ABN90, RRID: AB_11205592), IBA1 (Wako Chemicals Cat# 019-19741, RRID:AB_839504), CD68 (BioRad Cat# MCA1957, RRID: AB_322219); Neurexin 1 (Millipore Cat# ABN161, RRID: AB_10917110), and GFAP (Synaptic Systems Cat# 173004; RRID: AB_10641162). Fluorescent images were acquired using a laser scanning confocal microscope (LSM 800, Zeiss) or slide scanner (VS200, Olympus). All the images were blinded for genotype before subjecting to analysis using FIJI software from National Institute of Health.

### Neuronal Cultures and Immunocytochemistry

Primary cortical neurons from WT and CSPα KO mice (P0) were cultured on coverslips, as previously described (72). Immunocytochemistry was performed as done previously (72). Fluorescent images were collected with a Zeiss LSM 800 laser scanning confocal microscope (Oberkochen, Germany) with a 63× oil immersion objective. Primary antibodies used for this study are: Bassoon (Enzo Life Sciences Cat# ADI-VAM-PS003, RRID: AB_10618753), Homer 1 (Synaptic Systems Cat# 160003, RRID: AB_887730), CSPα (Chemicon, Cat# AB1576; RRID: AB_90794), ATG9a (Abcam Cat# ab108338; RRID: AB_10863880), Amphiphysin 2 (Millipore Cat#05-449, RRID: AB_309738).

### qPCR

RNA was extracted from CSPα KO and their WT littermate control frozen cortical tissues at P24 (n=6/group) utilizing the Qiagen RNA Mini Plus prep kit and recommended protocol. The quality of the purified RNA was measured using an Agilent bioanalyzer 2100. For real-time qPCR, cDNA was synthesized with the PrimeScript 1^st^ Strand kit with an additional primer mix containing random DTs (#RR047A and #6110A, Takara). cDNAs were amplified using specific Taqman probes, and iTaq Universal Probe Supermix and run on a C1000 Thermal Cycler (Bio-Rad) equipped with Bio-Rad CFX Manager software. A standard Taqman probe protocol was utilized: 95°C for 3:00 minutes; then cycles 40 times: 95°C for 0:10 seconds; 60°C for 0:30 seconds. The following Taqman probes (Thermofisher) were used: *Sept5* (Mm01175430_m1), *Dmn1* (Mm00802468_m1), *Actb* (Mm02619580_g1), *Gapdh* (Mm99999915_g1). Expression was determined by normalizing target expression to housekeeping genes (*Actb* and *Gapdh*) and corresponding CSPα KO values to the WT average and plotted using Prism 10.

### Single cell RNA velocity (scVelo) analysis and gene identification

RNA velocity was computed using the Python package scVelo. The CSPα KO and WT samples were processed separately to highlight the differences between these two conditions. First, the function filter_and_normalize was applied to conduct data filtering, normalization, and log transformation. Next, for each cell, the first- and second-order moments were calculated by the function pp.moments. Then, the gene velocities and graph were calculated, and the velocities were projected to the precomputed UMAP plot using velocity, velocity_graph, and velocity_embedding_stream functions respectively, with default parameters (38).

### CellChat analysis

The normalized data was first split into CSPα KO and WT samples for comparison analysis, then the CellChat objects were created by the function createCellChat, followed by subsetData. Subsequently, the over-expressed genes and ligand-receptor interactions associated with each cell group were identified using the functions identifyOverExpressedGenes and identifyOverExpressedInteractions. In addition, the communication probability and cellular communication network were inferred through computeCommunProb and computeCommunProbPathway. Finally, the aggregated cell-cell communication network was calculated by function aggregateNet (40).

### Quantification and statistical analysis

All analysis was done blind to genotype. A Student’s t-test with Welch’s correction was used to compare the two genotypes. The number of mice used for each experiment are described as ‘‘n’’ in Results and Figure legends. Values are expressed as mean ± standard error of the mean (SEM) and *p* value of 0.05 or less was considered statistically significant. We used GraphPad Prism (9.2.0) software to perform statistical analyses.

## Author contributions

N.W. and S.S.C. conceived and designed this study. N.W. performed single nucleus RNA sequencing experiments, genome alignment, single cell quality control, and clustering analysis. N.W. also performed ultrastructural analysis, immunohistochemistry, imaging, primary cortical neuron culture, western blotting as well as mouse genotyping. B.Z. performed computational analysis including DEGs, GO, RNA velocity, and CellchatDB analysis, with N.W. and S.S.C. providing guidance on biological interpretation, and H.Z. on statistical and computational methods. R.G. performed the qPCR experiment. N.W. and M.A. made the figures and illustrations. N.W., M.A., and S.S.C. wrote the manuscript. All authors have read and provided inputs to the manuscript.

## Acknowledgements

We thank Dr. Le Zhang and her lab member Haowei Wang for teaching us the nuclei isolation protocol. We would also like to thank Dr. Guilin Wang (Yale Center for Genome Analysis) for assistance with snRNA-seq, Dr. Xinran Liu & Kimberley Gibson (Yale Center for Cellular and Molecular Imaging) for assistance with electron microscopy, and Oliva Stebbins for quantifying a subset of electron micrographs. This work was supported by NIH grants 1R01 NS083846 (SSC) and 1R21NS132546 (SSC and NW) as well as the Nina Compagnon Hirshfield Parkinson’s Disease Research Fund (SSC).

---

**Figure S1 Phenotypes of P24 CSPα KO mice. A.** At P24, neuron density in the motor cortex of WT and CSPα KO mice was evaluated by immunostaining for NeuN (green), and visualizing DAPI (blue). CSPα staining (red) was used to confirm genotype. **B.** Quantification of neuron density for A. Note the NeuN positive cells were reduced in CSPα KO mice. Scale bar: 50 μm. n=5 brains/genotype. **C**. Dissociated CSPα KO neurons exhibit synapse loss. Synapses were labeled by antibodies to Bassoon and Homer in WT and CSPα KO cortical neurons at DIV14. Scale bar: 10 μm. Note partial overlap of Bassoon and Homer signals in WT (yellow; arrows) indicating juxtaposition of pre- and post synapses and their loss in CSPα KO. **D**. Quantification of synapse number in C, n=5 cultures/genotype. Data are shown as mean ± SEM **p < 0.01, t-test

**Figure S2 Cluster characterization of P24 mouse cortex. A**. Diagram of snRNA-seq pipeline. **B.** Bar graphs showing the nuclei number isolated per brain. **C.** Average number of nuclei isolated grouped by genotype. Mean ± SEM, n=3 brains/genotype. **D.** Bar graphs showing the number of genes sequenced per nuclei in each sample. **E**. Average number of genes sequenced per nuclei grouped by genotype. Mean ± SEM, n=3 brains/genotype. **F**. UMAP plot of WT and CSPα KO nuclei. Note high degree of overlap, indicating minimal batch effects after correction. **G**. Proportion of each brain cell type grouped by genotype. **H**. Dnajc5 expression level in each brain cell type. **I**. UMAP plot of mouse brain major cell types including 9 ExN and 5 InN subtypes. **J**. Heatmap showing expression of cell type-specific top marker genes in the subtypes of ExNs and InNs.

**Figure S3 Specific markers used for cluster characterization.** UMAP plots of snRNA-seq data of brain cortex showing cell-type-specific markers identifying each cluster. n = 89,464 total nuclei.

**Figure S4 Expression of heat shock and CSPα substrates genes in neurons. A.** Dot plot for the expression of heat shock genes in ExNs and InNs grouped by genotype. **B.** Western blotting of Hsc70 in synaptosomal lysates from WT and CSPα KO mice. CSPα and Tubulin were used as controls. **C.** Quantification of protein levels of CSPα and Hsc70 in CSPα KO samples relative to that of WT. N = 3 brains/genotype, Data are presented as mean ± SEM. *p < 0.05, t-test. **D.** Dot plot for the expression of genes encoding the substrates of CSPα in ExNs and InNs. **E.** qPCR for the gene expression of Sept5 and Dnm1 in WT and CSPα KO cortex. Data are normalized to ActB and GAPDH as housekeeping genes and represented as mean ± SEM; n=5-6 brains/genotype done in technical triplicates; ns, not significant, t-test.

**Figure S5 Glial cells show upregulated and downregulated DEGs. A, B, C, D.** Volcano plots for DEGs between WT and CSPα KO for (A) MG, (B) Astro, (C) Oligos and (D) OPCs.

**Figure S6 The interaction of glial cells and neurons. A.** Fan plot for the interaction strength of Astro and InNs. **B, C.** Heatmap of the increased receptor-ligand pairs between Oligo (B), OPC (C) and InNs in CSPα KO mice. **D, E, F, G.** Heatmap of the most differential receptor-ligand interaction pairs between Astro (D), MG (E), Oligo (F), and OPC (G) and ExNs. Scale: Communication probability of CSPα KO brains – communication probability of WT brains.

