## Supplementary figures and images for "Decoding transcriptomic signatures of Cysteine String Protein alpha-mediated synapse maintenance"

### Fig S1

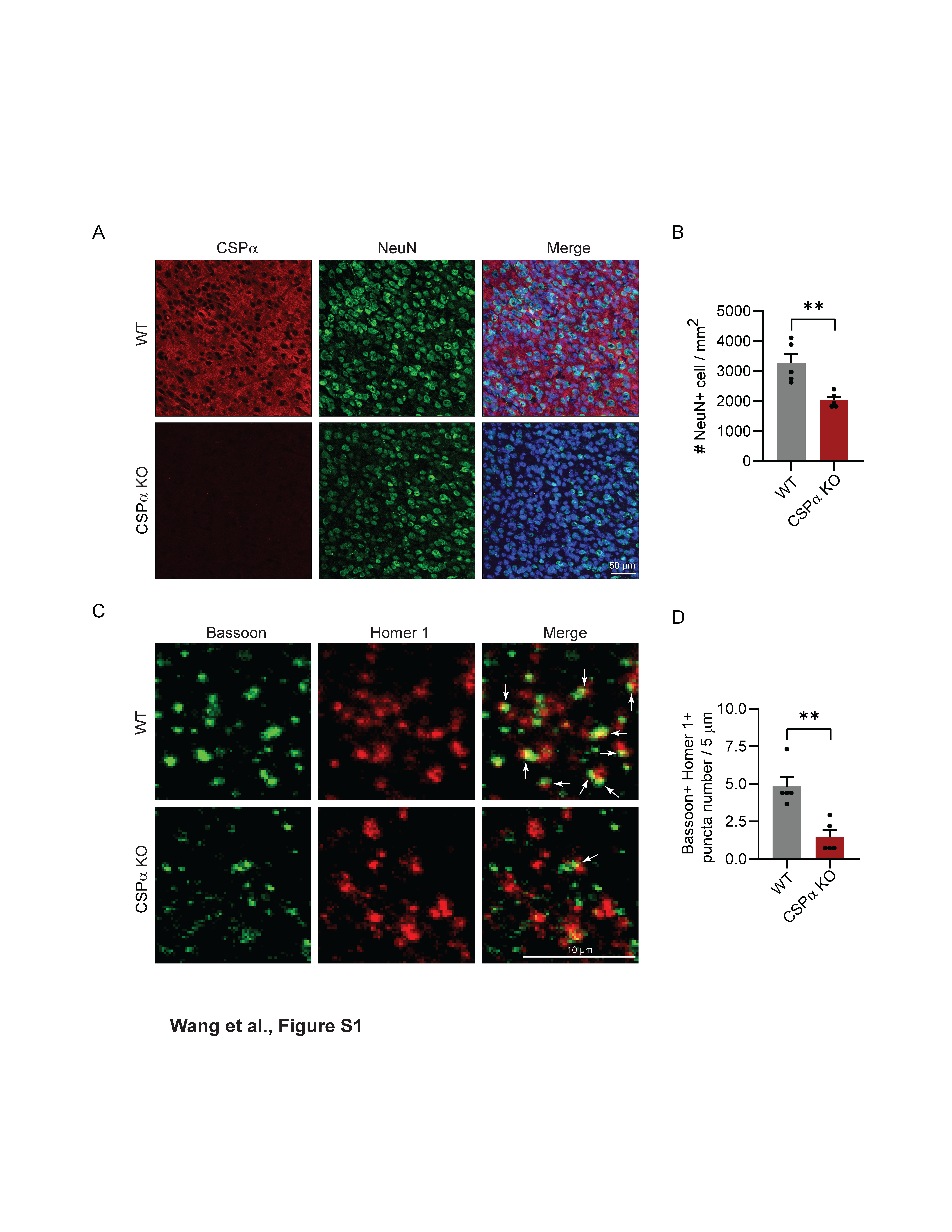

### Fig S2

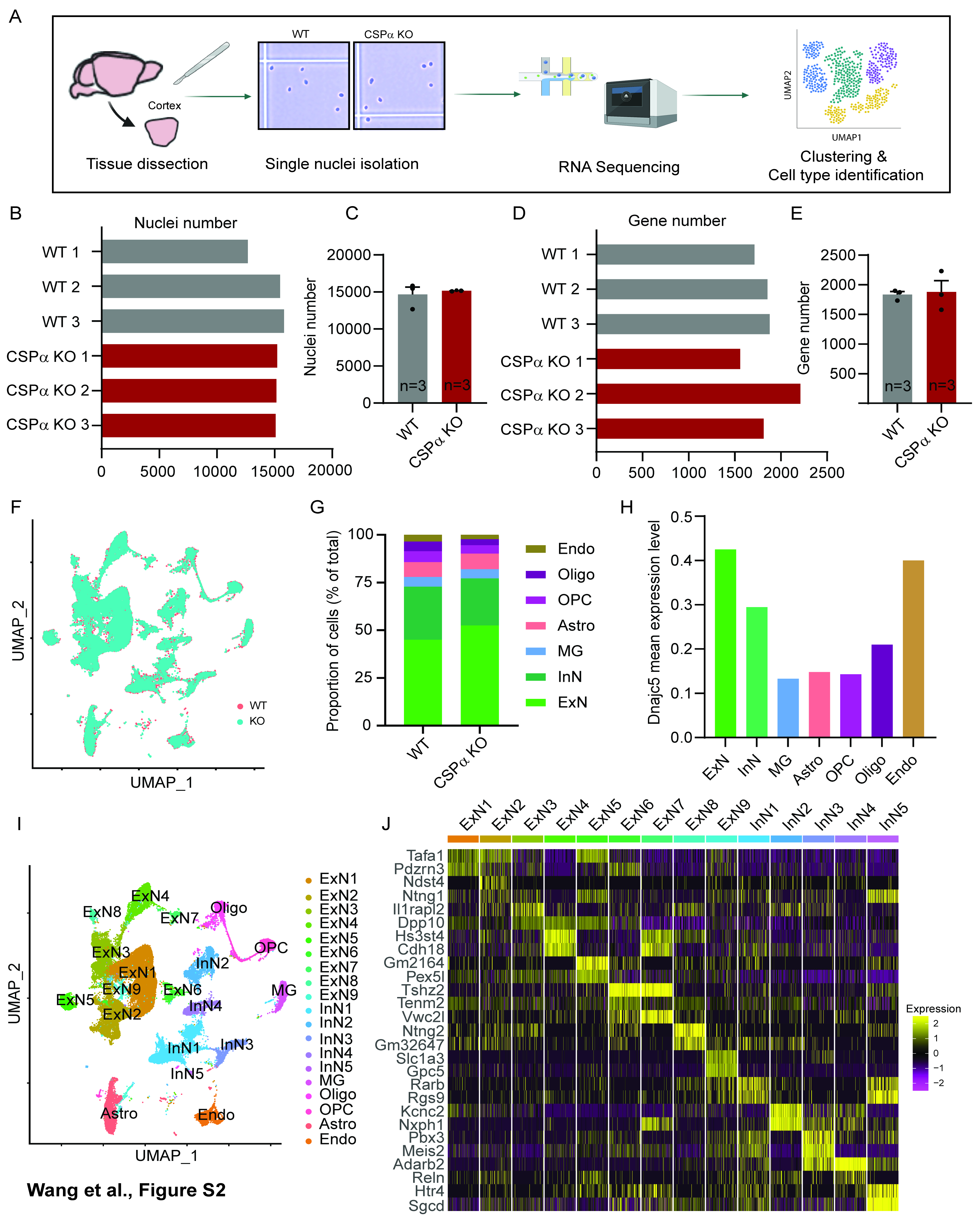

### Fig S3

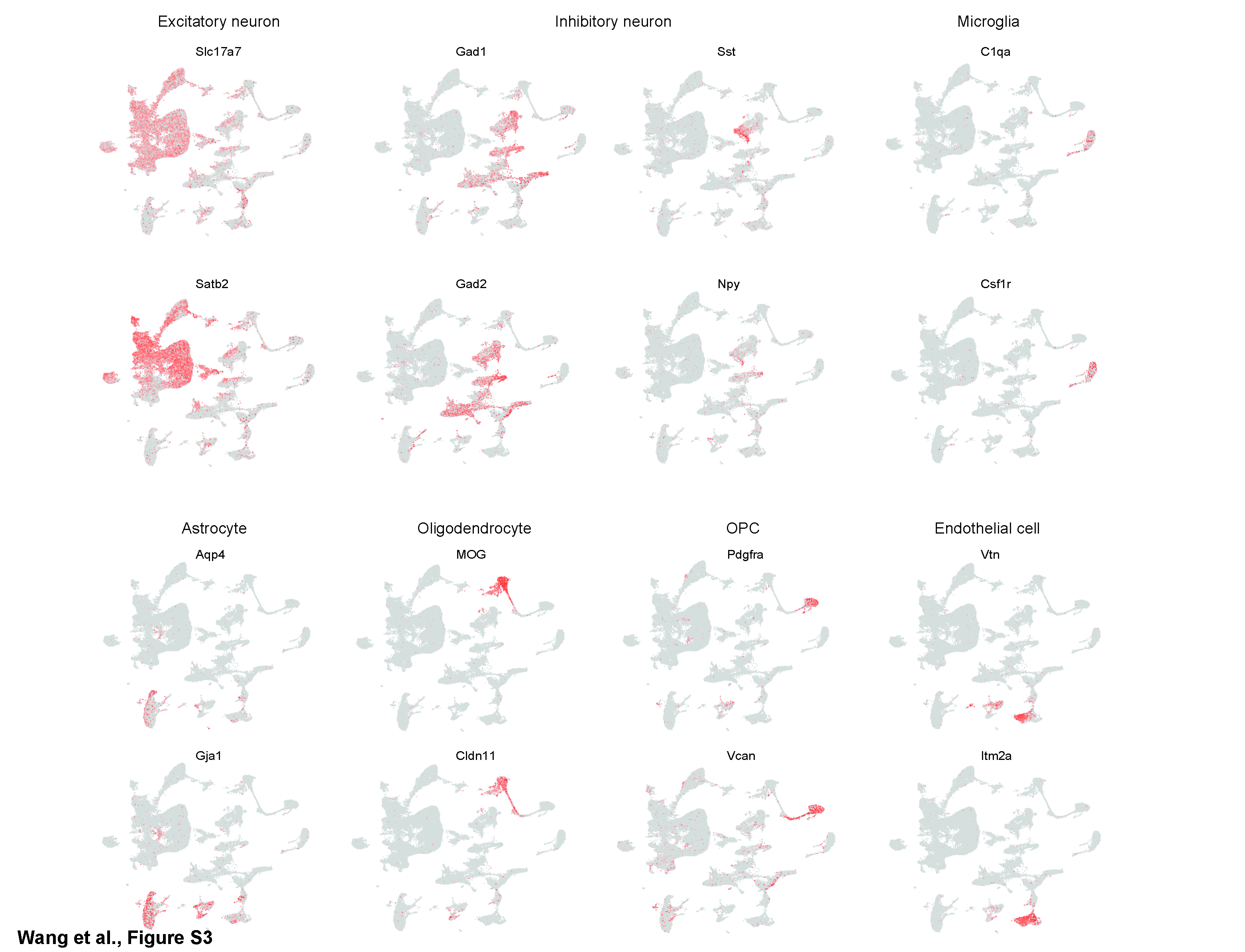

### Fig S4

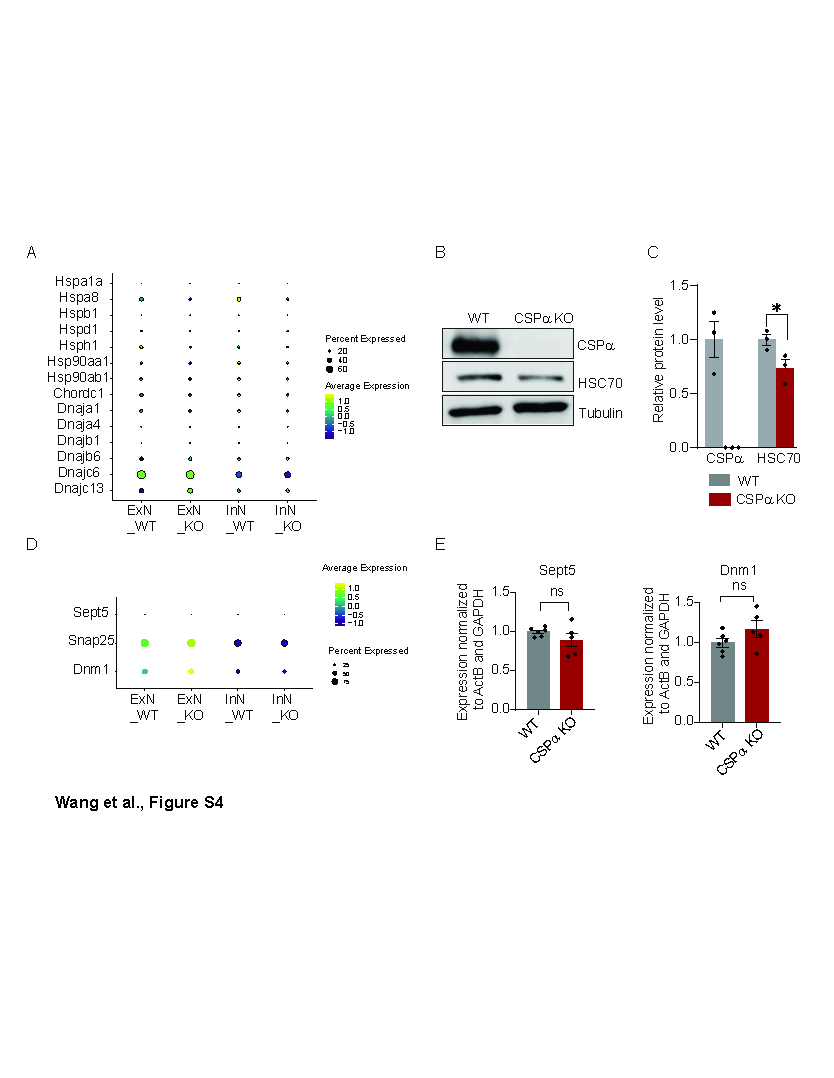

### Fig S5

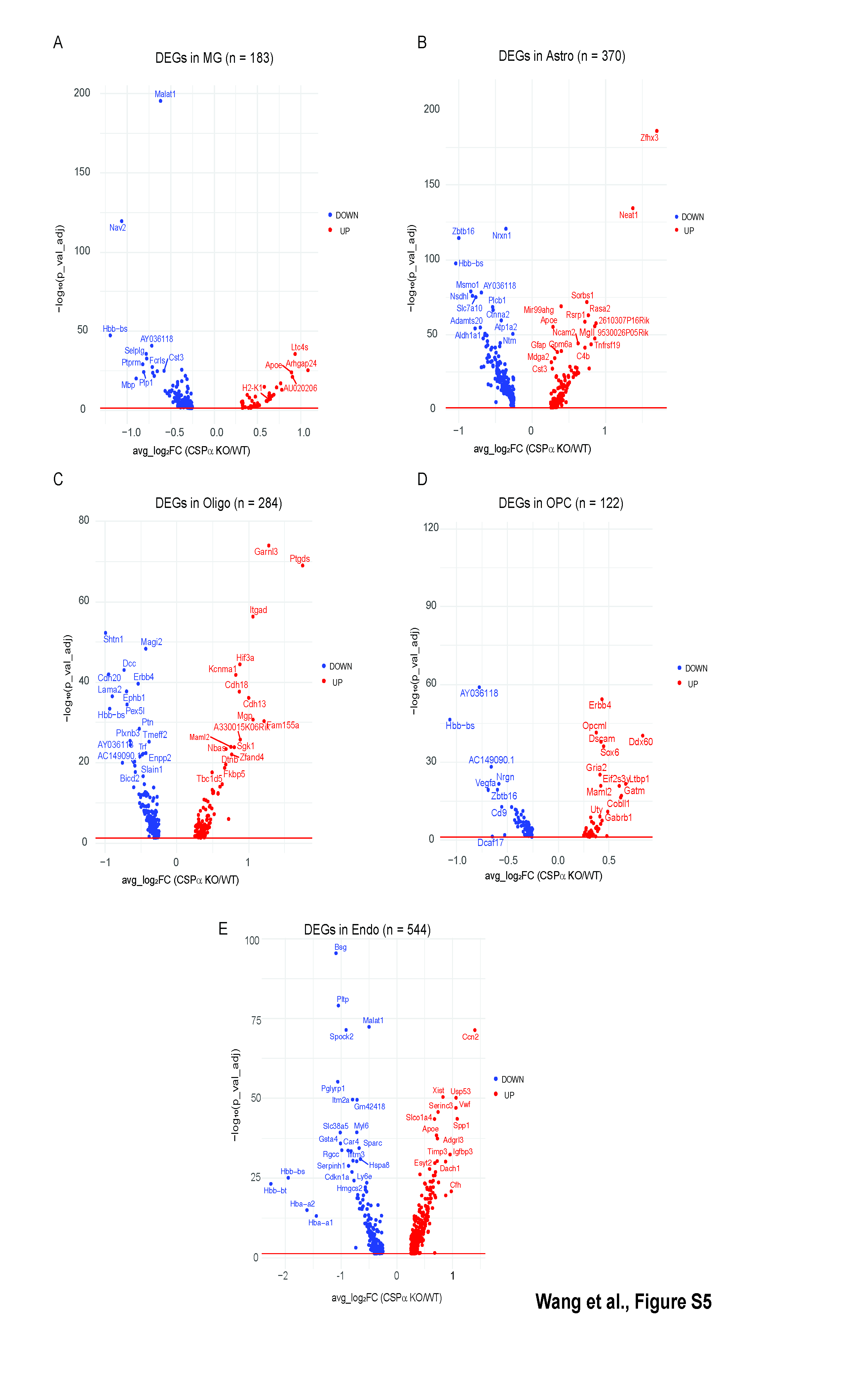

### Fig S6

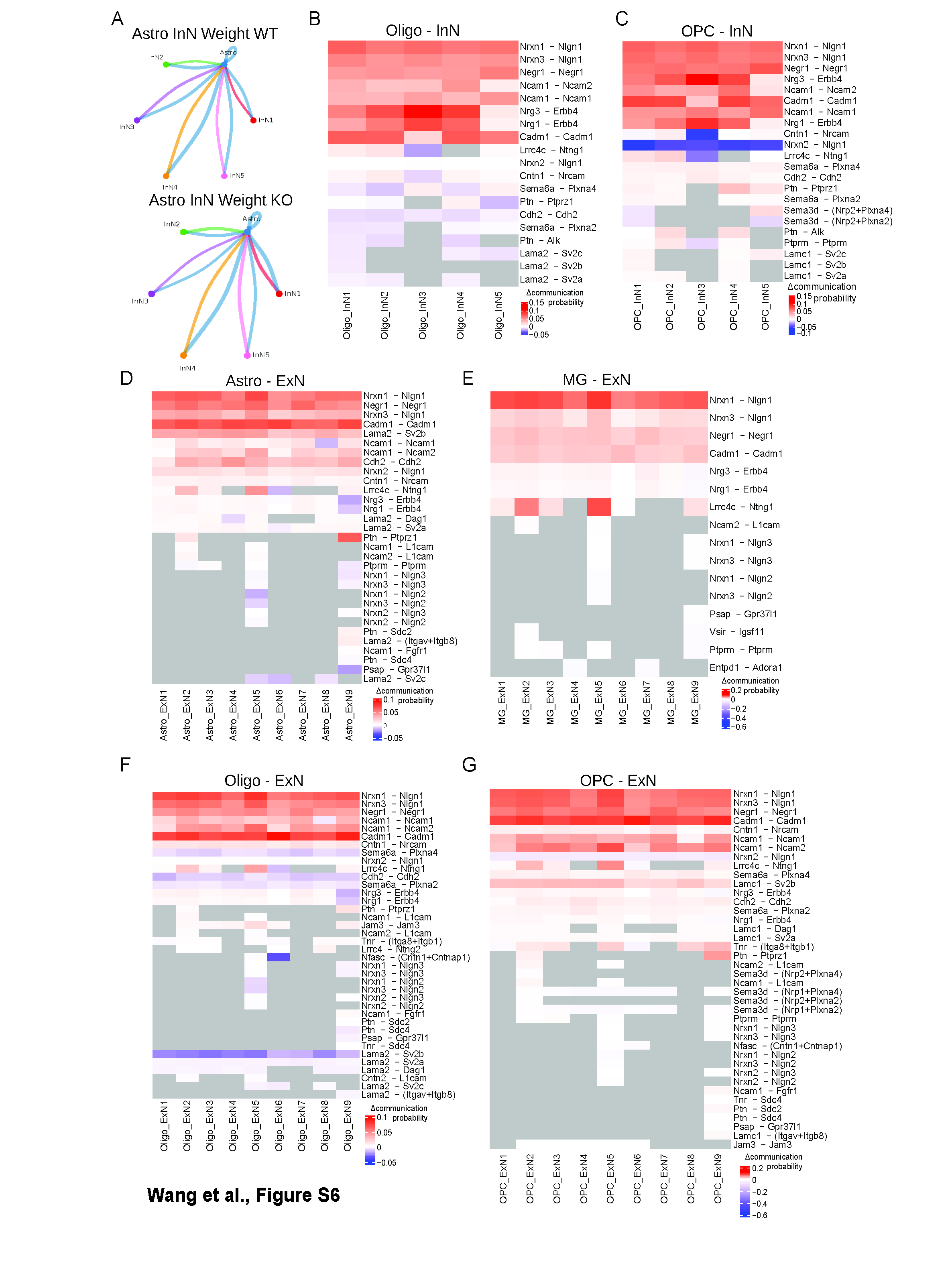
