## Supplementary material for "Decoding transcriptomic signatures of Cysteine String Protein alpha-mediated synapse maintenance": Table S1

| Cell Type | Pathway Name | Gene Name |
|---|---|---|
| ExN1 | Synapse organization | Asic2/Homer1/Ctnna2/Arc/Grid2/Atp2b2/Lingo2/Rph3a/Prrt1/Cacng2/Unc13a/Etv5/Ephb1/Dlg4/Sptbn2/Hspa8/Ntrk2/Gpc6/Dab2ip/Numb/Lrfn2/Lrrtm4/Sdk1/Bdnf |
| ExN2 | | Ctnna2/Homer1/Unc13a/Prrt1/Rph3a/Etv5/Rock2/Dlg4/Hspa8/Adgrl1/Shank1/Numb/Sptbn2/Ntrk2/Lrrk2 |
| ExN3 | | Asic2/Malat1/Homer1/Atp2b2/Arc/Rph3a/Cacng2/Nrxn3/Lrfn2/Lrp8/Unc13a/Ephb1/Etv5/Prrt1/Grid2/Nrxn2/Dlg4/Hspa8/Shank1/Sptbn2/Pak1/Nrp1/Bdnf |
| ExN4 | | Homer1/Nrxn3/Arc/Malat1/Unc13a/Pak1/Ephb1/Ctnna2/Nrg1/Etv5/Dlg4/Atp2b2/Asic2/Rph3a/Shank1/Adgrb1/Sptbn2/Nfasc/Prrt1/Clstn3/Igsf21/Cacng2/Rims4/Arhgap39/Ephb2/Ntrk2/Igsf9b/Srcin1 |
| ExN5 | | Atp2b2/Rph3a/Pak1/Cacna1a/Etv5/Ctnna2/Asic2/Prrt1/Homer1/Dlg4/Unc13a/Malat1/Sptbn2/Cacng2/Cabp1/Shank2/Nfasc/Rims4/Arhgap39/Sorbs1/Hspa8/Shank1/Ntrk2/Clstn3/Camk2b/Syngap1/Dgkz/Ephb2/Rock2/Tmem108/Myh10/Dip2a/Adgrb1/Sez6l/Gpc6/Ephb1/Igsf21 |
| ExN6 | | Malat1/Etv5/Asic2/Ctnna2/Unc13a/Ntrk2/Nrxn3/Homer1/Atp2b2/Cacna1a/Dab2ip/Grin2a/Arc/Sptbn2/Bdnf/Syndig1/Rock2/Shank1/Rph3a/Dlgap3/Syngap1/Hspa8/Shank2/Ephb2/Prrt1/Rims4/Dlg4/Adgrl1/Arhgap39/Lrrk2/Il1rapl1/Sez6l/Il1rapl2/Nrxn2/Epha4 |
| ExN7 | | Rph3a/Unc13a/Elfn1/Etv5/Atp2b2/Gpc6/Cacna1a/Homer1/Dlg4/Adgrl1 |
| ExN8 | | Asic2/Nrxn3/Malat1/Arc/Cacng2/Homer1/Nfasc/Unc13a/Lrfn2 |
| ExN9 | | Ctnna2/Arc/Homer1/Asic2/Rph3a/Atp2b2/Dlg4/Unc13a |
| ExN1 | Dendrite development | Ctnna2/Fat3/Arc/Map1b/Camk2a/Ephb1/Dlg4/Map6/Iqsec1/Srgap2/Kidins220/Ntrk2/Dab2ip/Robo1/Numb/Sdk1/Bdnf |
| ExN2 | | Ctnna2/Fat3/Map6/Camk2a/Rock2/Dlg4/Robo1/Shank1/Numb/Ntrk2/Kidins220/Lrrk2 |
| ExN3 | | Arc/Fat3/Iqsec1/Map6/Kidins220/Lrp8/Ephb1/Robo1/Dlg4/Shank1/Pak1/Nrp1/Fstl4/Bdnf |
| ExN4 | | Arc/Pak1/Robo1/Ephb1/Ctnna2/Nrg1/Dlg4/Shank1/Iqsec1/Map6/Iqgap1/Fat3/Fstl4/Ephb2/Ntrk2/Kidins220/Srcin1 |
| ExN5 | | Camk2a/Iqsec1/Pak1/Cacna1a/Ctnna2/Dlg4/Nedd4l/Shank2/Map6/Pacsin1/Shank1/Ntrk2/Srgap2/Camk2b/Syngap1/Ephb2/Rock2/Kidins220/Dip2a/Ephb1 |
| ExN6 | | Ctnna2/Ntrk2/Fat3/Camk2a/Cacna1a/Dab2ip/Arc/Map1b/Bdnf/Rock2/Shank1/Syngap1/Shank2/Kidins220/Ephb2/Dlg4/Nedd4l/Iqsec1/Map6/Lrrk2/Il1rapl1/Pacsin1/Epha4 |
| ExN1 | Regulation of synapse organization | Asic2/Homer1/Ctnna2/Arc/Grid2/Lingo2/Etv5/Ephb1/Hspa8/Ntrk2/Gpc6/Dab2ip/Lrfn2/Lrrtm4/Bdnf |
| ExN1 | Regulation of synapse structure | Asic2/Homer1/Ctnna2/Arc/Grid2/Lingo2/Etv5/Ephb1/Hspa8/Ntrk2/Gpc6/Dab2ip/Lrfn2/Lrrtm4/Bdnf |
| ExN3 | | Asic2/Malat1/Homer1/Arc/Lrfn2/Lrp8/Ephb1/Etv5/Grid2/Slc17a7/Hspa8/Bdnf |
| ExN6 | | Malat1/Etv5/Asic2/Ctnna2/Ntrk2/Homer1/Dab2ip/Arc/Bdnf/Syndig1/Syngap1/Hspa8/Shank2/Ephb2/Rims4/Adgrl1/Lrrk2/Il1rapl1/Il1rapl2/Epha4 |

| | | |
|---|---|---|
| ExN1 | Postsynapse organization | Homer1/Arc/Grid2/Rph3a/Etv5/Ephb1/Dlg4/Sptbn2/Hspa8/Numb/Lrfn2/Lrrtm4 |
| ExN2 | | Homer1/Rph3a/Etv5/Rock2/Dlg4/Hspa8/Shank1/Numb/Sptbn2/Lrrk2 |
| ExN3 | | Homer1/Arc/Rph3a/Lrfn2/Lrp8/Ephb1/Etv5/Grid2/Nrxn2/Dlg4/Hspa8/Shank1/Sptbn2/Nrp1 |
| ExN4 | | Homer1/Arc/Ephb1/Etv5/Dlg4/Rph3a/Shank1/Adgrb1/Sptbn2/Arhgap39/Ephb2/Srcin1 |
| ExN5 | | Rph3a/Etv5/Homer1/Dlg4/Sptbn2/Shank2/Arhgap39/Sorbs1/Hspa8/Shank1/Camk2b/Syngap1/Dgkz/Ephb2/Rock2/Tmem108/Myh10/Dip2a/Adgrb1/Ephb1 |
| ExN6 | | Etv5/Homer1/Grin2a/Arc/Sptbn2/Rock2/Shank1/Rph3a/Syngap1/Hspa8/Shank2/Ephb2/Dlg4/Arhgap39/Lrrk2/Il1rapl1/Nrxn2/Epha4 |
