## Supplementary material for "Decoding transcriptomic signatures of Cysteine String Protein alpha-mediated synapse maintenance": Table S2

| Cell Type | Pathway Name | Gene Name |
|---|---|---|
| InN2 | Synapse organization | Ctnna2/Malat1/Rims4/Unc13a/Hspa8/Etv5/Rock2/Homer1 |
| InN3 | | Malat1/Prkca/Homer1/Atp2b2/Sybu/Camk2b/Cntnap4/Shank2/Dock10/Lrrtm4/Abl2/Asic2/Cdkl5/Cttnbp2/Hspa8/Ntrk2/Tiam1/Unc13a/Nrcam/Ank3/Numb/Epha3/Syngap1/Slc1a1/Cacna1a/Grin2a/Nrg2/Syn1/Dlgap3/Ntng1/Tnr |
| InN4 | | Rock2/Ntrk2/Rims4/Il1rapl1/Hspa8/Unc13a/Lrp8/Nrg2/Rapgef4 |
| InN2 | Asymmetric cell division | Zbtb16/Etv5/Sox5 |
| InN2 | Regulation of synapse organization | Ctnna2/Malat1/Rims4/Hspa8/Etv5/Homer1 |
| InN4 | | Ntrk2/Rims4/Il1rapl1/Hspa8/Lrp8/Nrg2/Rapgef4 |
| InN2 | Regulation of synapse structure or activity | Ctnna2/Malat1/Rims4/Hspa8/Etv5/Homer1 |
| InN2 | Regulation of cation transmembrane transport | Kcnc1/Stac2/Plp1/Ubash3b/Homer1/Cemip |
